# Brain, music and emotion: An EEG proof-of-concept study on musically continuous, non-personalized emotional responses

**DOI:** 10.1101/790972

**Authors:** Efthymios Papatzikis, Anri Herbst

**Affiliations:** Faculty of Communication, Arts and Sciences, Canadian University Dubai, Dubai, U.A.E.; South African College of Music, University of Cape Town, Cape Town, South Africa

## Abstract

It has been repeatedly reported that motivation for listening to music is majorly driven by the latter’s emotional effect. There is a relative opposition to this approach, however, suggesting that music does not elicit true emotions. Counteracting this notion, contemporary research studies indicate that listeners do respond affectively to music providing a scientific basis in differentially approaching and registering affective responses to music as of their behavioral or biological states. Nevertheless, no studies exist that combine the behavioral and neuroscientific research domains, offering a cross-referenced neuropsychological outcome, based on a non-personalized approach specifically using a continuous response methodology with ecologically valid musical stimuli for both research domains. Our study, trying to fill this void for the first time, discusses a relevant proof-of-concept protocol, and presents the technical outline on how to multimodally measure elicited responses on evoked emotional responses when listening to music. Specifically, we showcase how we measure the structural music elements as they vary from the beginning to the end within two different compositions, suggesting how and why to analyze and compare standardized, non-personalized behavioral to electroencephalographic data. Reporting our preliminary findings based on this protocol, we focus on the electroencephalographic data collected from n=13 participants in two separate studies (i.e., different equipment and sample background), cross-referencing and cross-validating the biological side of the protocol’s structure. Our findings suggest (a) that all participants – irrespectively of the study – reacted consistently in terms of hemispheric lateralization for each stimulus (i.e., uniform intra-subjective emotional reaction; non-statistically significant differentiation in individual variability) and (b) that diverse patterns of biological representations emerge for each stimulus between the subjects in the two studies (variable inter-subjective emotional reaction; statistically significant differentiation in group variability) pointing towards exogenous to the measurements process factors. We conclude discussing further steps and implications of our protocol approach.

## Introduction

Research studies indicate that listeners respond affectively to music, often associating basic or primary emotions such as happiness, sadness, fear and anger with musical stimuli [for example, 1-20]. Konečni [21–23] adds the sublime as a further category of evoked responses, while it has also been found that listeners clearly react to music’s specific structural elements (according to the level of their musical expertise [24] to form a final emotionally charged evoked response [25, 26]. It must be noted that any emotional association with or evoked response to music starts early in life. Even three-year-old children are capable of associating musical excerpts with emotions [27, 28].

Although a scientific basis for approaching and registering affective responses to music^i^ mostly exists with specific focus on either their behavioral or biological states separately (for example the MRI study by Koelsch *et al.* [29]; or Küssner & Koelsch [30]), it is rare – if existing at all – to find studies that have combined the behavioral and neuroscientific research domains, providing a cross-referenced neuropsychological and non-personalised outcome that successfully connects standardized semantic emotion labels with personal perception to biology, *specifically* using a continuous response methodology with ecologically valid musical stimuli of the *same* cohort for both research domains. It is even rarer for studies to follow the aforementioned path using specifically the EEG modality in their experimental investigations [10, 11, 31–36].

## Background

In providing a larger context for this study, Altenmüller *et al.* [11] used cortical direct current electroencephalography (dc-EEG; focusing on bands between 0.01-1Hz) to measure 160 short complex sound sequences comprised of 15sec excerpts; 120 musical and 40 environmental ones. At the behavioral level, they measured the short sound sequences using a Likert scale which ranged from (1) “like very much” to a (5) “do not like at all,” matching personalized semantic labels to the related evoked responses. Although the experiment was entirely successful in its parts, it did not use extended and continuous – hence ecologically valid – musical stimuli. Ιt also did not manage to fill the *non-personalised semantic code-personal perception* gap as it did not measure emotion but rather preference.

Sammler *et al.* [31] on the other hand, intentionally manufactured short consonant (happy) and dissonant (sad) organized non-musical sound sequences (1 min long) yet to avoid the personalization issue mentioned above. In their study, they managed to measure and report objectively on the Alpha band activation correlation to the valence distinguished evoked responses (happy-sad) in an eyes-closed condition. However, as they did not follow a musicologically elaborate example (musical composition), an ecologically valid musical stimulus was not fully evident in this case too, projecting a limited framework of generalization in music-specific neuropsychologically defined evoked emotions.

On the contrary, the EEG approach that took place in the Mikutta *et al.* studies [33, 34] investigated possible correlates of experience-driven changes of emotional arousal, presenting a continuous and ecologically valid stimulus (Beethoven’s 5^th^ symphony first movement; 7.4 mins) to both amateur and professional musicians. The researchers examined the tonic and phasic sides of the electrophysiological data they collected, finding that experienced musicians rate their physiological arousal in a more consistent way than amateur musicians, which is on par with their elevated brain activation. However, the researchers did not use a standardized platform of registered emotional ratings to the music stimulus, providing in the end only subjective arousal measurements. These measurements brought to the fore an intrapersonal dimension of evoked responses to the ecologically valid musical stimulus, however as a rather subjective perceptual construct of a musical experience, that led to a personalized emotional reaction as opposed to an objective reaction. One could therefore speculate if it was the level of the musical experience (i.e., attention to specific details or lack thereof; knowledge or lack of certain musical information) that influenced musicians to rate their emotional responses in a consistent or inconsistent way, or if it was the actual content of the musical piece being known to them (i.e., anticipation and knowledge of expectancy) that led to the final result of consistency. It furthermore raises the question of whether unfamiliarity or incapacity to understand the piece would lead to a different result? Would the emotional result – hence, brain activation – and rating be similarly consistent?

Sourina, Liu, and Nguyen [32] worked on more novel research designs, slightly filling the gap of standardized comparative measurements between behavior semantics and biology. As in the previous case, however, the major part of their work focused on the personalized dimension of explaining musically evoked emotions rather the study of an actual non-personalised correlation between the behavioral semantics of emotions and their parallel brain activation. Using a real-time EEG-based human emotion recognition algorithm, the research team incorporated fractal dimension values extracted from the available EEG-Sound envelops cross-registration, pinpointing indeed a detailed path on the EEG analysis using a mixture of a standardised pool of affective sound stimuli – the International Affective Digital Sounds (IADS) [37] as well as 1 min long non-standardised musical excerpts (i.e., the standardized material was not ecologically valid, while the ecologically valid material was not standardized). The behavioral measurements, although following a non-personalised standardization path, were not cross-referenced to the neuroimaging data, failing to address the inter-modularly assigned semantic labels. In addition, only six basic emotion labels were used, excluding a broad spectrum of available emotional labels, therefore most probably ‘disguising’ potentially stated evoked responses and perceived emotional states.

Finally, the Hsu *et al.* [35] recent study followed a similar approach as Sourina *et al*. with adding an extra layer of data and therefore better understanding towards the ‘semantics-perception-biology’ system discussed here. More specifically, the researchers focused on devising a loop protocol, bidirectionally feeding the studied dimensions of information (behavior and EEG), while managing to present a ‘transformational matrix,’ based on an algorithm which created a personalized ‘semantics-perception-biology’ framework. Although the researchers bidirectionally fed and successfully aligned the two investigated dimensions, it is of our opinion that in this case too, the EEG cross-correlation to the behavioral semantics was not very strong, as (a) the team did not fully employ an ecologically valid musical stimulus (they used the IADS as in the aforementioned study), hence limiting the emotional perception and measured response, while (b) they based the whole experiment on the subjects’ real-time self-feeding and tempering of the model’s data – through a well-developed UI – that most probably interacted with the overall projected emotional content and its registration without offering any control for this component of the data.

In summary, the above studies show a growing attempt to refine combining standardized measurements of all three neuropsychology and musicology domains (behavior-biology-musical structure). However, it is also evident that an all-inclusive protocol, where neuropsychological and musicological methods are employed equally, has not been presented so far. In this sense, all aforementioned research protocols may have left aside valuable information that musicology with specific reference to musical structure has to offer in understanding emotionally-evoked responses in the brain and behavior, and whether the musical structure (as a whole – e.g., measurement and time series of a musical piece – or in parts – e.g., a cadenza, appoggiaturas or chord progression) may interact with and alter the neuropsychological system in a standardized way, regardless of the level of the musical experience. This, at least, has already been found to be only the case for the behavioral side of things [38, 39].

## Aims and Hypothesis

Considering all the above, the aim of our research project is to measure for the first time in a multimodal way emotional responses to structural elements as they vary from the beginning to the end in a composition (variation measured through continuous response throughout the musical composition) within different compositions, analyzing and comparing standardized behavioral to electroencephalographic data. Our general research hypothesis is that emotion, projecting a particular *valence or intensity to a specific (or a set of) structural element(s) of a musical stimulus, provides a joint behavioral and electroencephalographic synchronization imprint* to all participants when registered in an ecologically valid testing environment.

Following the complexity of the aim and our hypothesis, as well as to be more transparent and concise in the way we explain our findings and *Modus Operandi*, we decided to present the essential components of our protocol in separate parts – as proof of concept written constructs – making known in detail the process and practical implications that such an endeavor entails. Therefore, in this article, we do not aim to present the final joint findings and analysis of both the behavioral and neuroimaging data, rather provide an in-depth discussion of the conceptual framework of our EEG protocol. A brief presentation of the behavioral part is, however, included.

## Research Method

For the behavioral part, in brief, we followed a grounded theory approach [40] by looking for emergent themes when analysing a combination of various tasks (drawings [41, 42], self-report narratives and choosing of adjectives from the updated standardized Hevner list (see [43]). For the electrophysiological part – the main focus of this article – we based our work on the Schmidt and Trainor [10] model (see further below for more information), focusing on the EEG data collection and analysis stage for a defined sample (n=13), aiming to replicate the electrophysiological process that Schmidt and Trainor followed in their experiment. Compared to all other related and previously discussed experiments, our study’s novelty was that we, first of all, run two mini-studies (Study 1 and Study 2) with different equipment aiming towards a cross-verification of results. Furthermore, we created and used for the musical stimuli a non-personalized (standardized) measurements setting, applied to both the electrophysiological and behavioral approaches.

A brief outline of the combined neuropsychological research protocol follows (see also Fig 1):

1. Recording of participants’ neurophysiological baseline (ABR-DPR);
2. Presentation of two different musical stimuli and neurophysiological recording of emotional responses. The two musical stimuli were segregated by white noise;
3. Repetition of Step 1;
4. Presentation of the two previous musical stimuli and behavioral registration of emotional responses through drawings, written narratives and choices of emotional responses from a provided adjective list;
5. Individual interviews to clarify, decode, and describe responses in Step 4.

**Fig 1.**
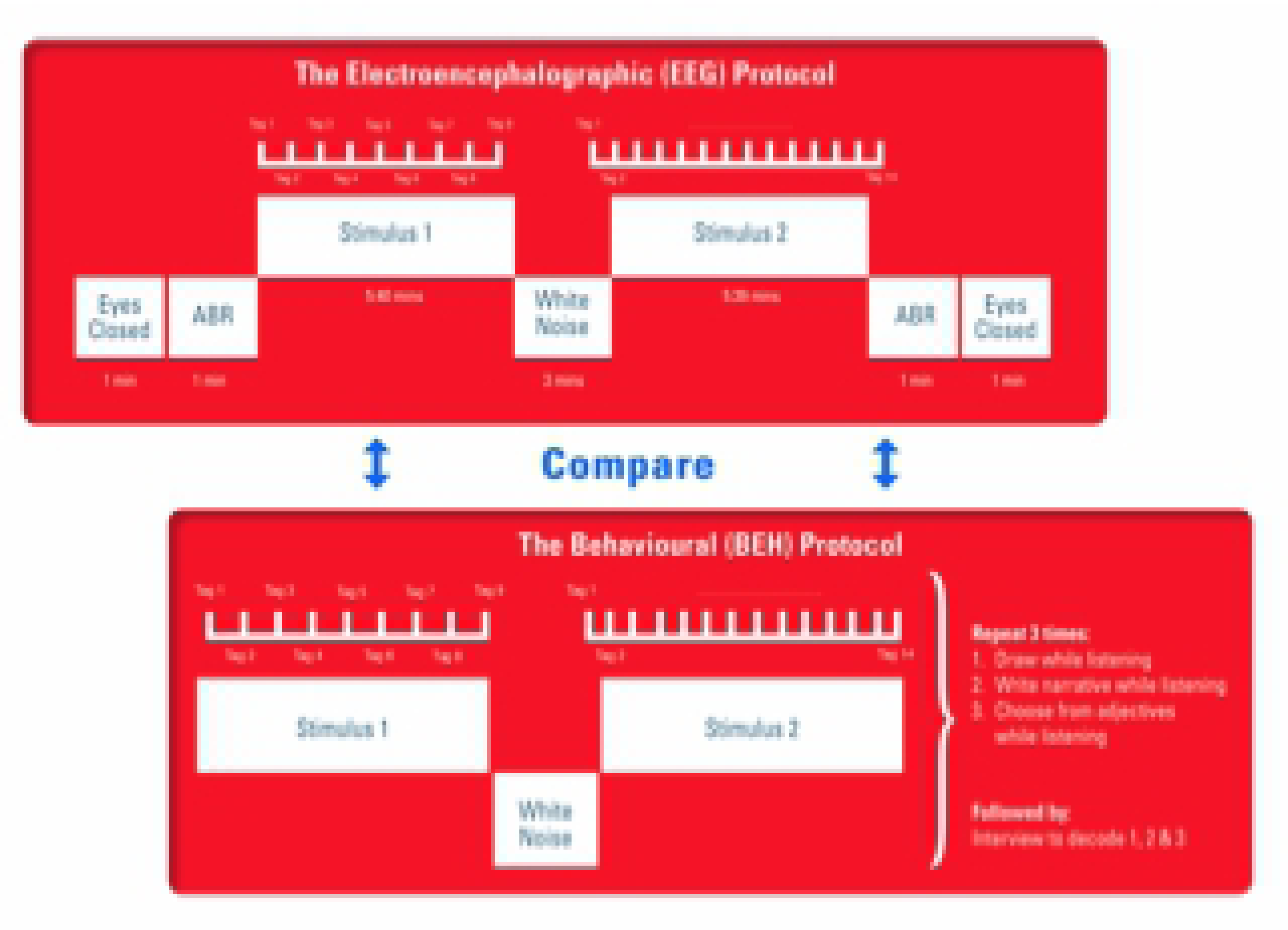
The EEG and Behavioural Protocols Combined

As far as our data analysis method is concerned, our statistical computations followed a similar binary analysis approach for both measured modalities (behavior and brain data), calculating repeated measures (RM) ANOVAs and independent ANOVAs for the within- and the between-subjects conditions respectively, according to a predetermined emotion-musical-structure related framework (musicological aspect). For the EEG part, more specifically, we focused on the hemispheric lateralization of activity (dependent variable) and followed the alpha synchronization paradigm. This paradigm has been shown to denominate qualitative states of emotional valence [10, 11, 44, 45].

## Materials and procedures for the EEG

### The Music Stimuli

The following two solo violin compositions were accordingly prepared in WAV-lossless formats and used as musical stimuli to collect our data:

(Composition A) Nikos Skalkottas (1904–1949), Sonate for Solo Violin, fourth movement: Adagio – Allegro molto moderato (1925). Duration: 5”40’. Performed by Georgios Demertzis.

(Composition B) Hendrik Hofmeyr (b. 1957), *Luamerava* for Solo Violin, Commissioned in 2000 by the South African Music Rights Organisation. Duration: 5’39”. Performed by Ian Smit.

We chose these two solo violin works because they are similar in genre, timbre (instrument) and duration, as well as being unfamiliar to most, if not all participants. They furthermore met the criteria of not containing lyrics and being composed in two distinct countries, one North of the equator and one South.

The South African composition by Hendrik Hofmeyr was composed post-apartheid and the Greek Skalkottas composition after the First World War. Both composers lived outside of their home countries in Europe for extended periods. Hofmeyr spent ten years in Italy and Skalkottas spent 12 years in Germany. Hofmeyr is outspoken about his strong links with tonality, and his composition styles could be described as representing neo-Classical and neo-Romantic sound ideals [46–48]. On the contrary, Skalkottas was influenced by Schönberg’s serialism, and he held high regard for Bartók [49–50].

The Hofmeyr work has a philosophical basis on the sonification of the music tonality of the Shona *Mbira dza vadzimu* with its associated overtones that the composer combines with his compositional ideas. *Luamerava* is furthermore loosely based on Mutwa Credo’s work “*Indaba my Children*” and his reference to an African goddess [51]. Although not strictly programmatic in nature, this composition is undergirded by extra-musical ideas, which might be reflected in the participants’ responses. In contrast, the Skalkottas composition is idiomatic at every turn, perhaps even more so than the later Sonata for Solo Violin by Bela Bartók. The Solo Sonata by Skalkottas refers to Baroque models, combining the Greek meta-war folk musical structure, and an atonal system that the composer often follows in his works. This work provides an extensive background of musical emotions in linear and complex forms, thus showcasing a depth of content to study and discuss. The choice of these two musical stimuli holds much potential for the study of affect, as it could render very different participant responses.

Each of the two musical compositions was approached as a continuous sound stimulus, hence a hypothetical emotional macro-response. However, since knowing that specific structural elements (e.g., tempi, chord progressions, modulations, pitch ranges, cadenzas, instrument ranges, playing techniques) can also evoke emotional micro-responses in a musical composition [25, 26] – all resulting in the relevant macro-response – we also decided to identify the micro-responses in our study. To achieve this, we asked a systematic musicologist^ii^ to perform a music-theoretical analysis. This analysis was cross-referenced and verified by two other professional musicologists, based on aura-based and score-based analyses as well as a combination of two.

The music-theoretical analysis identified and converted to time-specific tag points the micro-responses of potential emotional change and produced noted memos in the musical scores presented in Western staff notation. Most importantly, this binary micro- and macro-emotional response approach enabled us to pinpoint and capture inter-subjective as well as intra-subjective differentiations or similarities of neuronal oscillations for each musical composition in the specified timeframes, hence to study emotional responses both at a micro-as well as a macro-neurophysiological time-related-to-emotion frame.

The analysis of the Skalkotas composition rendered nine specified timepoints that we refer to as tag points. The Hofmeyr rendered 14. These tag points for each composition are shown in Table 1, presenting the corresponding time duration in minutes and seconds since the beginning of each musical piece. Based on these tag points, we identified the independent and dependent variables of the experiment. The i*ndependent variables* were our participants and the two musical compositions they listened to, whereas the *dependent variables* were the hemispheric lateralization of the alpha band activation, as well as the lateralized brain activation effect for and between each tag point.

**Table 1.**
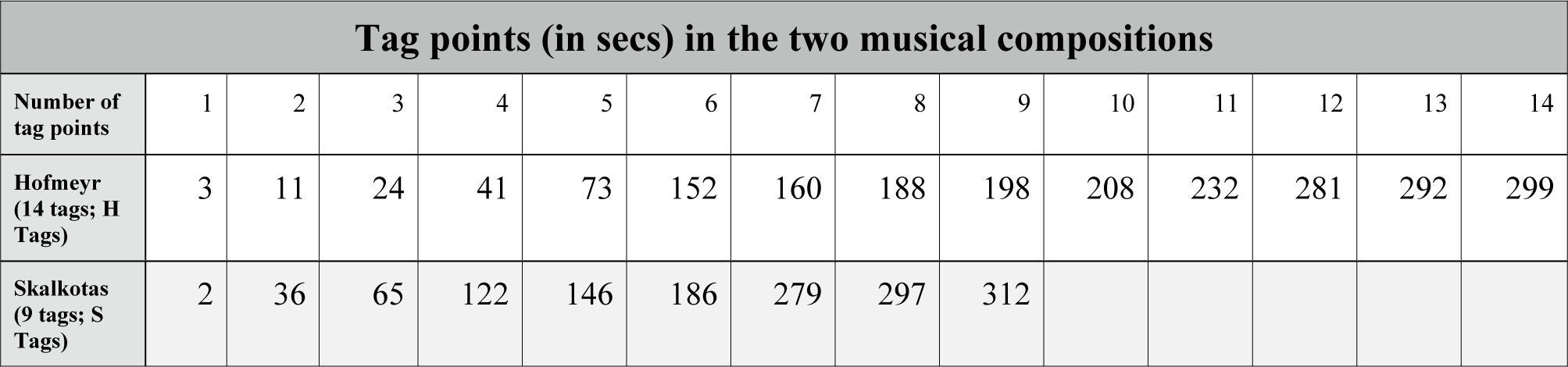
Tag points distribution

| Tag points (in secs) in the two musical compositions |  |  |  |  |  |  |  |  |  |  |  |  |  |  |
| --- | --- | --- | --- | --- | --- | --- | --- | --- | --- | --- | --- | --- | --- | --- |
| Number of tag points | 1 | 2 | 3 | 4 | 5 | 6 | 7 | 8 | 9 | 10 | 11 | 12 | 13 | 14 |
| Hofmeyr (14 tags; H Tags) | 3 | 11 | 24 | 41 | 73 | 152 | 160 | 188 | 198 | 208 | 232 | 281 | 292 | 299 |

| Tag points (in secs) in the two musical compositions |  |  |  |  |  |  |  |  |  |
| --- | --- | --- | --- | --- | --- | --- | --- | --- | --- |
| Skalkotas<br>(9 tags; S<br>Tags) | 2 | 36 | 65 | 122 | 146 | 186 | 279 | 297 | 312 |

### Participants

#### Study 1

Three participants (music experts) (n=3) were randomly recruited from the Music Department, the South African College of Music of the University of Cape Town (UCT). They were attending the university for three or more years as music students, and took one of the two following courses amongst playing their instruments and taking other music courses: Music Theory and Analysis III/IV and Music History of Western music III/IV. All three subjects played a musical instrument on a third-year level. Testing on the subjects took place after Ethics Clearance following the Helsinki declaration was obtained from the Human Research Ethics committee at the Faculty of Health Science, UCT. Participants took part in the project once they have given full written informed consent, and only after they received detailed information during a relevant presentation pertaining to what is required of them and how the experiment works. They were informed about their right to withdraw at any time they see fit. All three right-handed participants had no mental health or drug usage history, while no physical hearing injuries or abnormalities were shown during the experiment. Their ages were 23, 24, and 30 (Mean=25.7). At the time of the experiment, they were enrolled for BMus degrees at UCT and were in their final year of study. All three continued with postgraduate studies: Honours in Orchestral Conducting, Masters in Music Therapy, and Masters in Musicology. As it has been shown that perception and expression of emotions are differently imprinted in the brain for right-handed (RH) participants versus the left-handed (LH) participants [52, 53], we recruited right-handed subjects to avoid possible data analysis complications and distortions.

#### Study 2

For the second study, thirteen participants (n=13) were randomly recruited from the Metropolitan College, Thessaloniki, Greece. All of them were related to music through formal music education (instrumental and theoretical backgrounds), having variable studies range extending between 2 and 10+ years. Although the majority had some music tuition in their youth, not all studied music at tertiary level. Some were professional musicians or students at the College at various stages in their studies. This is a significant difference between Studies 1 and 2, but not detrimental to the experiment as it is important to measure if the number for years to formal music instruction would play a significant role in the findings. No mental health or drug usage history existed before the experiment, while no physical hearing injuries or abnormalities were shown during the experiment. Testing took place only after the Ethics process was concluded according to the steps we followed in Study 1, including the Metropolitan College research administration’s ethical approval in this case as was further required. For Study 2, the mean age of the final 10 participants we successfully processed, was 26.3 (Mean), while out of the 13 initial participants, the remainder were mostly right-handed (7RH, 3LH). Regarding the three excluded participants, one had to voluntarily withdraw from the EEG recording phase (personal reasons), while the other two could not be finally processed due to the very noisy raw data obtained from them (i.e., artifacts mostly generated because of excessive movement).

### The Schmidt and Trainor Study

The Schmidt and Trainor study, a pioneer in its field, aimed at two distinctive goals. The first was “to examine whether different musical excerpts induce different affective states that can be indexed by measuring brain activity” [10], while the second was “to examine whether measures of regional EEG activity can be used to differentiate emotions along the two dimensions of valence and intensity simultaneously” [10]. The whole study was critically based on three specific regional brain activation and emotion models: firstly, primarily combining emotion to brain activity [44]; secondly, concerning the role of absolute frontal activation in the intensity of emotion [54, 55], and thirdly considering hemispherically lateralised brain activity in relation to emotional valence and intensity [45].

In summary, the Schmidt and Trainor study investigated the relationship between regional brain activity evoked by music listening and induced emotional responses. The researchers attempted to directly relate valence and intensity of emotional experience to electrical activation in the brain through EEG while presenting to a group of undergraduates (n=59) orchestral musical excerpts designed to induce joy, happiness, fear, and sadness. The four musical excerpts were pre-rated by a different group of undergraduate students to represent “intense-unpleasant (fear), intense-pleasant (joy), calm-pleasant (happy) and calm-unpleasant (sad)” [10].

The investigators used the four orchestral excerpts (60secs/each) to record the participants’ EEG activity, placing four electrodes on the left and right mid-frontal (F3, F4) and parietal (P3, P4) scalp regions, according to the 10/20 Electrode Placement System [56]. All electrodes were referenced to the Cz region, while the collected raw data were bandpass filtered between 1Hz and 100Hz, and sampled at 512Hz. Electrooculography (EOG) was also performed to facilitate further deartifacting. The raw data were visually inspected to exclude eye blinks, eye movements, and other motor movements, using specialized software by James Long Company (EEG Analysis Program, Caroga Lake, NY). Finally, the artifact-free data were analyzed using a Discrete Fourier Transform, while plotting results were outputted in the alpha band between 8Hz and 13Hz.

Among the combined 2×2 ANOVA computations on gender, valence (pleasant-unpleasant), intensity (intense, calm) and hemispheric lateralization (right, left) that were performed in the study by Schmidt and Trainor, the valence by hemisphere analysis presented significant effects, providing a significant interaction between valence and hemisphere. Clear hemispheric lateralization was observed for the four musical excerpts in relation to their nominated emotion, projecting higher activity on the left hemisphere for positive emotions, and in reverse, higher activity on the right hemisphere for negative emotions.

### The EEG Data Collection

An electroencephalogram detects and registers the brain waves or electrical activity of the brain. During the registration procedure, the EEG electrodes detect tiny electrical voltages that result from the activity of the brain cells. The voltages are then amplified through specific equipment (the amplifier, a computer, and a software program) and appear as a graph on the screen.

#### Study 1

EEG data were collected using a 128-channel EGI Geodesic Sensor Net (GSN) system, found in the Division of Biomedical Engineering, University of Cape Town. EGI-GSN does not require any scalp preparation or abrasion, making it a very convenient choice due to its ease of application on many different subjects in a short period. The system’s combined temporal-topographic data collection abilities furthermore allow for a more detailed topographic signal analysis due to the close intersensor distance.

The electrodes were placed following the EGI GSN128 electrode placement system, while referenced to the central vertex (Cz). Data were collected through the following five steps:

1. Eyes Closed (EC) 1-minute session: This session took place in order to calculate the Posterior Dominant Rhythm (PDR) of all subjects. It has been found that the presence of a normal PDR indicates a well-functioning brain. A normal PDR is framed between 8 Hz and 12 Hz and is usually found at the posterior parts of the brain (e.g., O1-O2 electrode regions) during an eyes-closed condition. A proper PDR may rule out all sorts of pathology. If found to be less than 8.5 Hz or greater than 11.5 Hz, this should be considered abnormal in adolescents and adults [57, 58].
2. Auditory Brainstem Response (ABR) 1-minute session: An Auditory Brainstem Response measurement took place in order to estimate hearing sensitivity objectively in the participants. The ABR can be perceived as similar in the manufacturing process to the Auditory Steady-State Response (ASSR) [59] although the former’s electrical oscillations, hence the measurement reference, occur as a reaction much earlier in the time-brain region domain (i.e., brainstem versus neocortex). The ABR can be thought of as an electrophysiological response of the brainstem to rapid auditory stimuli. Its basic stimulus structure is usually comprised of clicks or tone bursts. ABR relies on statistical calculations to determine if and when a particular threshold is present. In our study, we used the following stimuli configurations to measure the ABR: two conditions of broadband spectral square wave clicks of 500Hz and 4000Hz lasting 30s each. One thousand sweeps of clicks were included in each one of the two 30 secs segments (500Hz and 4000Hz), each having a duration of 100 μsecs, and an inter-stimulus interval (ISI) rate of 33.3Hz. The two conditions were manufactured at an air conduction binaural acoustical level in the range of −14.1 and −14.3 dBFS, as shown in the following Fig 2. A 51 msecs ISI separated the two conditions.
3. Presentation of Compositions A and B segregated by White Noise (A-WN-B). Both stimuli were acquired and used in a WAV-Lossless format, while delivered through the Shure SE315 in-ear sound module. The Shure SE315 uses a vented balanced armature driver. These in-earphones have a frequency range of 22Hz–18Hz and their sound isolating design blocks out most ambient noise. Before using the musical stimuli, they were both normalized and equalized to achieve the best possible sound quality. Following the protocol of Eerola *et al*. [60], the music stimuli were equalized in terms of loudness, using peak RMS (root mean square) value normalization and careful subjective evaluation. The process of normalization can be defined as the boosting of “the highest sample of a digital sound file to the maximum amplitude the system is capable of encoding, short of CLIPPING (0dBFS), and then raising all other samples by the same proportion, in effect, raising the volume of the sound file, so it is as loud as possible without going into overload. This process maximizes resolution and minimizes certain types of noise” [61]. The entire A-WN-B set was played through, following the same room conditions (light, temperature, air ventilation) for all participants. EEG was continuously recorded following an eyes-open registration approach. Previous EEG studies [62–64] investigating the difference between the eyes-open versus the eyes-closed recording condition have shown that there is a difference in the power level of the final raw data registration, and that the eyes-open approach should be preferred, as the opposite presents a high-risk raw data reading result in the whole scalp EEG registration process [63]. Additionally, the Schmidt and Trainor study [10] had been conducted following an eyes-open registration process, too.
4. The final two sessions repeated the first and second steps for purposes of cross-referencing and evaluation of intrapersonal data.

**Fig 2.**
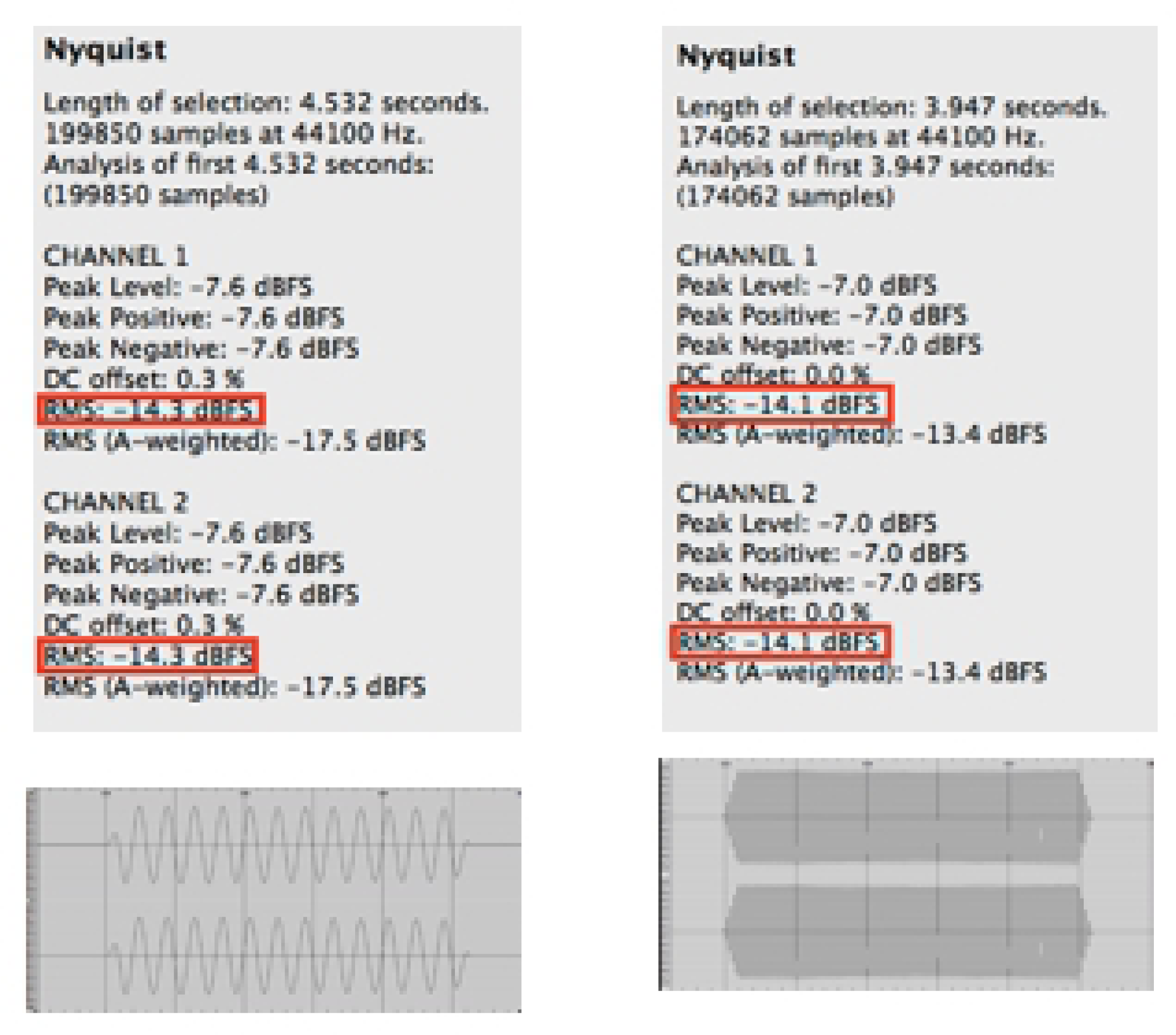
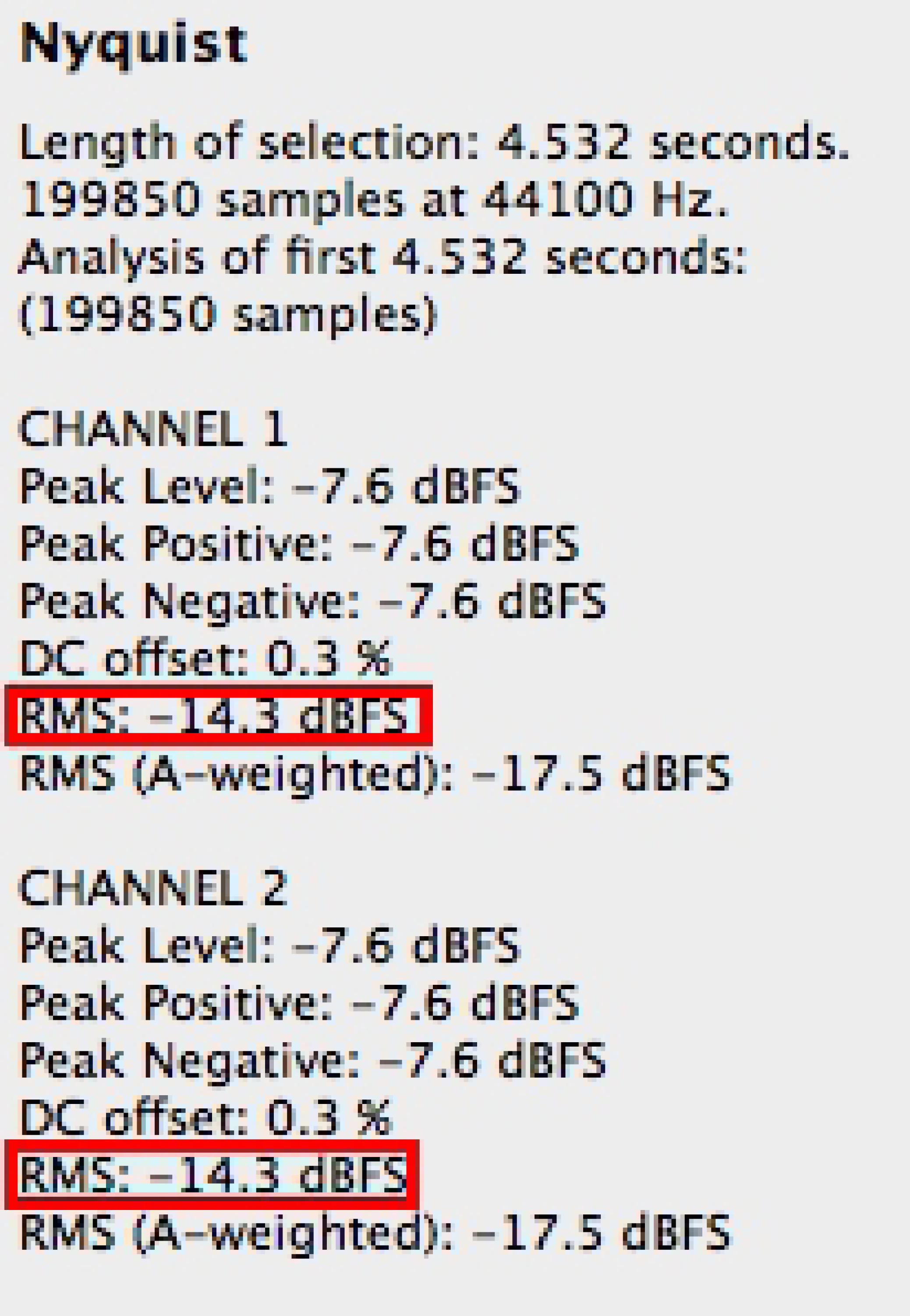
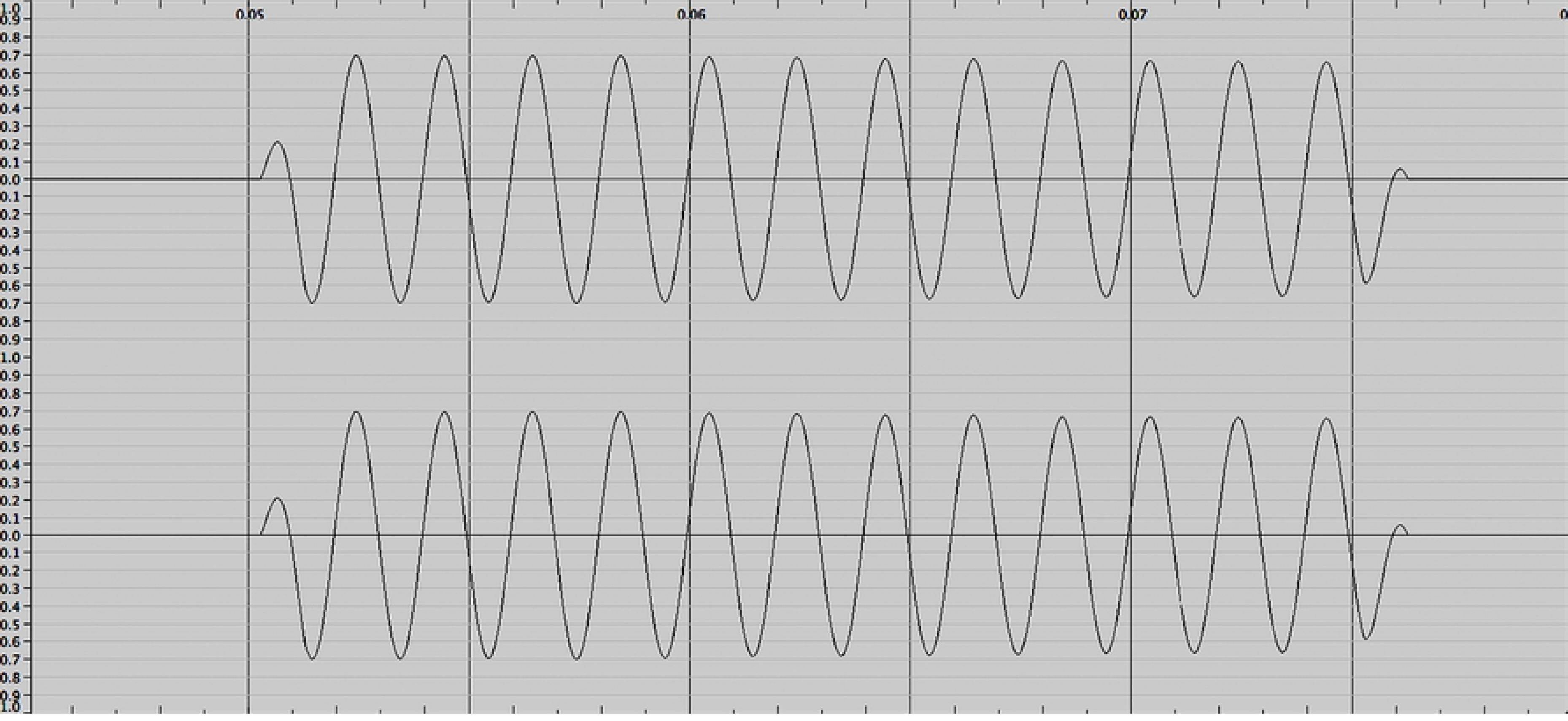
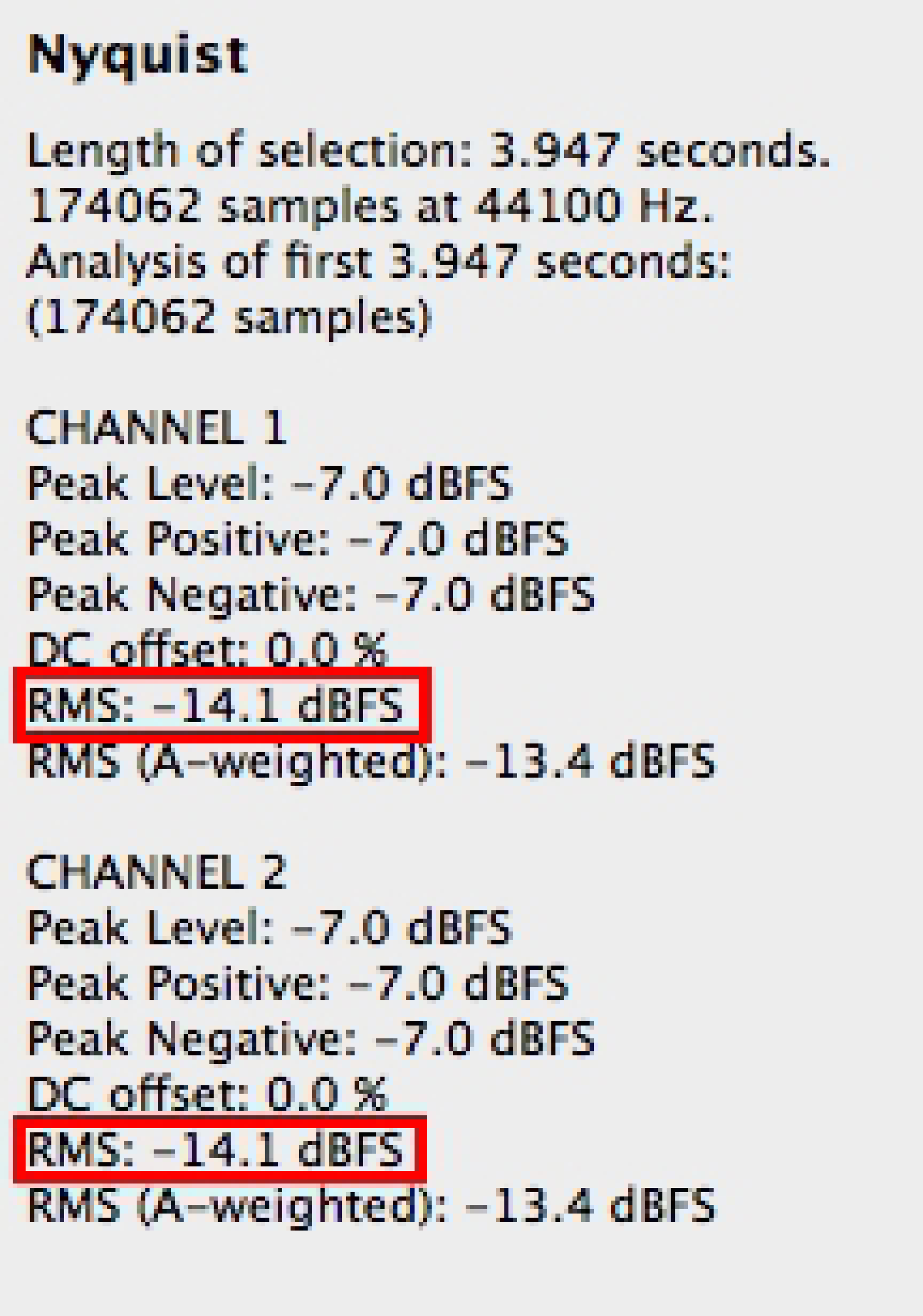
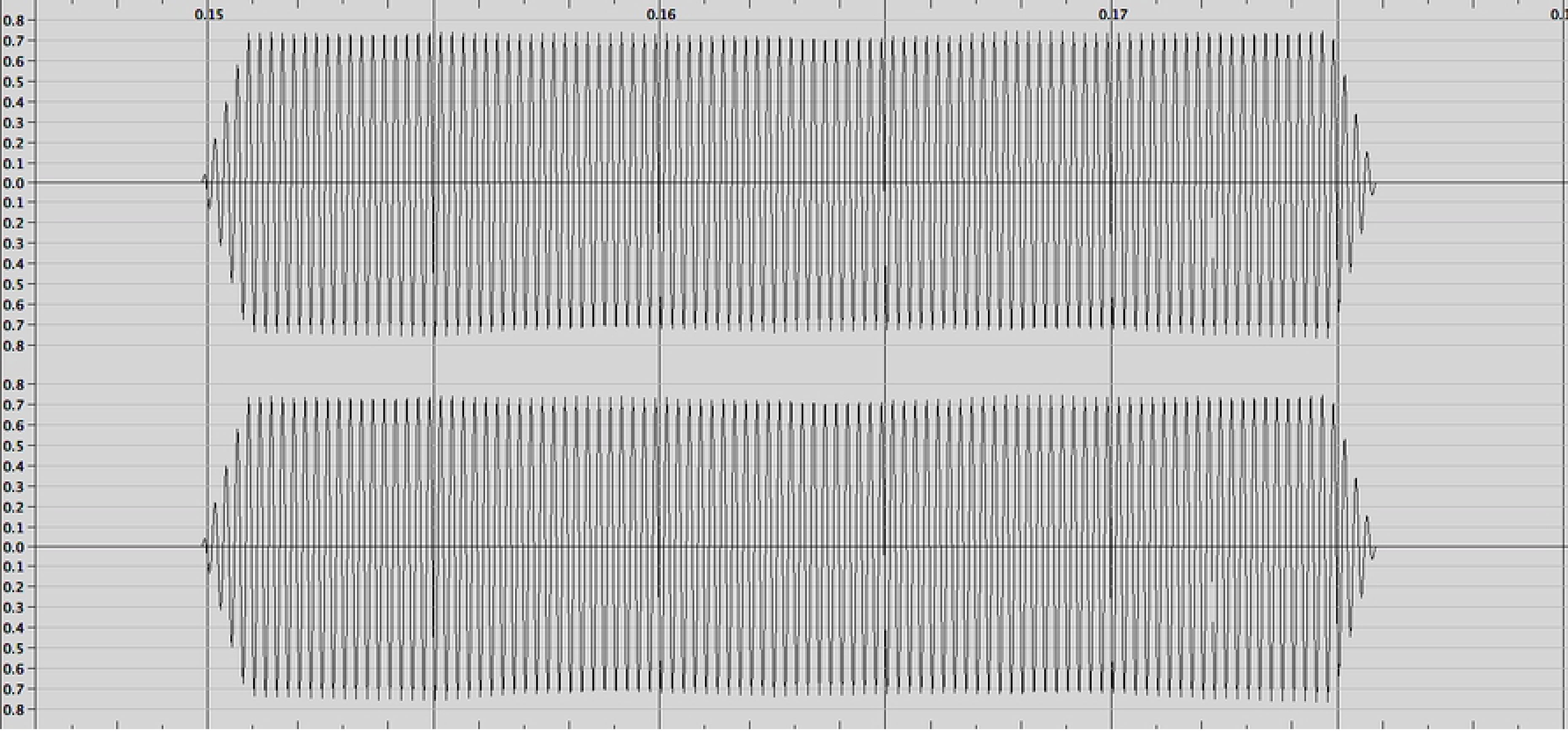
The ABR stimuli. The ABR stimuli were created using the PureData and ProTools software platforms. The signal polarity was following a full spectral wave circle (positive to negative alternatively) showcasing a 0 to positive initial deflection. To ascertain the RMS (Root Mean Square) of the ABR sections, about 4–5 seconds of each condition was initially selected, and an audio statistical gathering plugin was then applied in Audacity, using Nyquist to get the above information.

#### Study 2

EEG data were collected using the Emotiv EPOC+ 14 high-resolution channel system. The Emotiv has saline-soaked felt pads and does not require any scalp preparation or abrasion. It is a very convenient unit to use, yet not as sensitive as the EGI Geodesic Sensor Net (GSN) we used in Study 1. For the Emotiv system, the electrodes (AF3, F7, F3, FC5, T7, P7, O1, O2, P8, T8, FC6, F4, F8, AF4) were placed in accordance to the International 10-20 electrode placement system, while they were referenced to the CMS/DRL electrodes placed on the mastoids. Data were collected through the same four steps as presented above for Study 1.

### Why EEG and not MRI

It has been found that in humans, emotional recognition related to music occurs just after the first 500ms of the onset stimulus [65]. This fact implies that the process of emotional alteration can potentially be extremely fast and abrupt, while the effect of the musical stimulus onset and offset may greatly vary across narrow time frames [66], involving multiple cortical activations and de-activations, as well as the interaction of multiple neural networks. Based on this, identifying a suitable brain-imaging tool to measure emotional reactions to music can be challenging. The two standard tools usually employed in most of the relevant research protocols (EEG and MRI) each carry different strengths and weaknesses [67, 68].

As indicated before, our experiment aims to measure multimodal ways in which a group of musicians responds to structural elements within two different musical compositions. We, therefore, had to focus on the temporal rather than the spatial domain of collecting neuroscientific data, as the structural elements inducing different emotions within the different music compositions may greatly vary across narrow timeframes and raw data sound epochs. Facing this dilemma, EEG proved more suitable than fMRI for the following reasons:

Firstly: EEG is more precise in temporal data measurements than the fMRI. The MRI technique is not known to measure direct neuronal activation, but rather an indirect effect of this activation as represented in the oxygen ratio of the existing hemoglobin in the blood flow at a particular brain region (oxyhemoglobin and deoxyhemoglobin ratio). This effect is called the Blood Oxygen Level Depended (BOLD), with which researchers gain a window on the temporal dynamics of the brain hemodynamic response.

Nevertheless, it is also known that “the BOLD response [even for a very short stimulation period; approximately 1 second in duration] is typically delayed by 1 to 2 seconds and reaches its maximum after approximately 8s (typically between 5 and 10 s)” [68]. Therefore, there is always a relevant delay in the recording of data related to the temporal domain during the fMRI scanning process, which makes it very difficult to collect and analyze condensed information or rapidly changing evoked responses. On the contrary, the EEG module, by measuring the electrical activity of the brain directly, without delay, can achieve a detailed recording of changes in brain activation, targeting even the first milliseconds of a specific stimulus onset or offset [67]. In our case, using EEG assisted us in investigating all relevant emotional changes of selected structural elements in the chosen musical stimuli.

Secondly: As aforementioned, emotional responses to music may involve multiple cortical activations and de-activations, as well as the interaction of multiple neural networks. One result of these complex interactions is the presence of differential responses across brain regions using the fMRI approach, may lead to incomplete modeling of the relevant evoked bio-emotional elements, as we will not be able to follow a straightforward application of standard general linear model (GLM) approaches that previous fMRI/emotion research projects have used (for a brief overview, please see, for example [16]). Therefore we will not be able to perform a detailed comparison of the tomographic data that our research project will show with already existing ones, because of a non-uniform starting point of emotional/temporal reference. Previous fMRI research projects have only referred to relevantly more significant and ‘thinner’ in quality (happiness; sadness) blocks of evoked emotional responses through music, while our project endeavors to investigate thick, both in time and quality, alterations of emotions, connecting abstract emotional responses (i.e., drawings, narrative expressions) to biological representations in the brain. We believe that by using initially the EEG module in our project, and later on the sLORETA software [69], we will be able to study the electromagnetic tomography in relation to the evoked emotional responses and therefore more efficiently cross-reference it with essential studies in the field – the Schmidt and Trainor study in our case – in order to validate it.

Thirdly: Using the EEG module in our study instead of the MRI module, avoids the statistical interaction effect between the MRI scanner noise and the affective processes. It has been strongly suggested (Skouras, Critchley & Koelsch, 2013) that the MRI scanner noise can influence and distort affective brain processes like the ones we would like to investigate.

## EEG data processing and analysis

### EEG processing

In order to process and analyze the EEG raw data for both studies, we used the MatLab (Mathworks co.) EEG Lab version 11 software platform. The data were first montaged and segregated into epochs, following first an ‘as-per-condition’ block segregation and then an ‘as-per-tag-points’ segregation. A graphical illustration has been already presented in our protocol design earlier on (Fig 1). The following EEG epochs were created: ‘Eyes Closed,’ ‘ABR,’ ‘First musical stimulus’ (with pinned tag points), ‘White Noise,’ ‘Second musical stimulus’ (with pinned tag points), ‘ABR,’ ‘Eyes Closed.’ These epochs were deartifacted using both a manual (i.e., thresholding, by eye trimming) as well as a logarithmic signal decomposition approach (i.e., Independent Component Analysis/ICA).

In more detail, the following configurations/steps were applied in creating and processing these epochs:

1. Whole EEG data baseline removed – initially sampled in 250Hz;
2. Average montage for all electrodes;
3. Data were filtered using a 1Hz low-pass and a 50Hz high-pass filter;
4. False channels/data interpolated when needed;
5. ICA through the ADJUST algorithm performed;
6. Detailed artifact rejection performed;
7. ‘Epoching’ at tag-point level;
8. Hemispheric lateralization categorization (Left-Right dominant hemispheric activation; only for Study 1, please see explanation below);
9. Major dipoles calculation (only for Study 1, please see explanation below);
10. Computation of alpha-band asymmetry ratios on the closest to the four ‘Schmidt and Trainor’ electrodes. For Study 1, these were electrode numbers 25, 124, 53, and 87 [71]; for Study 2, these were electrodes F3, F4, P7, P8.

Specifically, for the epoching process at the tag-point level (step 7 above; similar process for Studies 1 & 2), the expectant points of emotional alteration (i.e. tag points) were noted on the EEG timeline axis according to the available musicological analysis producing specific EEG data envelopes (1sec pre-trigger stimulus up to 2secs post-trigger stimulus; see Fig 3). These data envelopes were fully extracted from the whole EEG data set for a separate tomographical and statistical analysis as described below.

**Fig 3.**
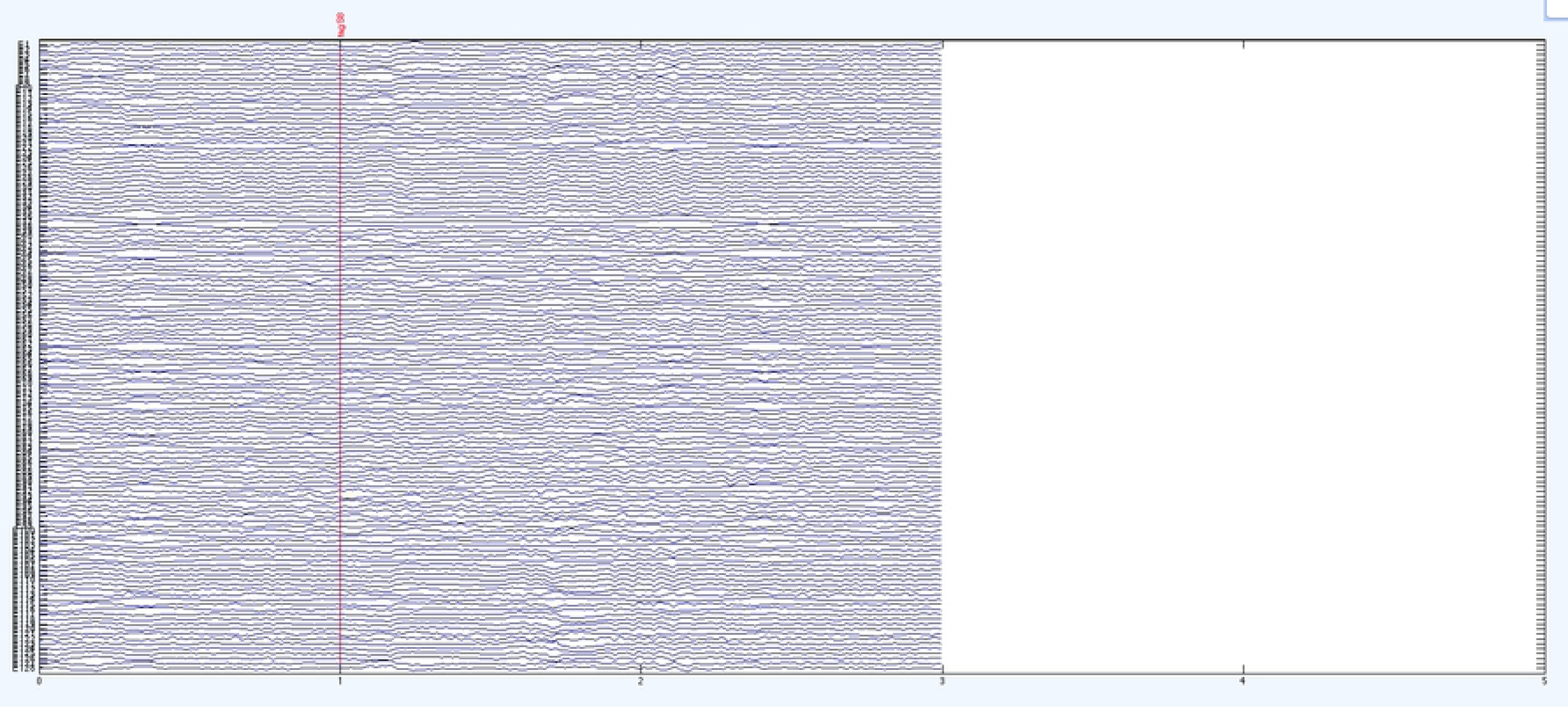
Example of extracted tag epoch; Subject 3 Tag S8 from Study 1.

### Analysis Design - Hemispheric Lateralization

In order to study the hemispheric lateralization dependent variable, we focused on the created epochs (step 7 mentioned above). On these epochs, we followed three steps of analysis for Study 1 (steps 8, 9, and 10) while only one step (step 10) for Study 2. The reason for not following the same detailed path for both studies was that statistical insignificance was found on the mean difference between the two sets of measurements (whole set of electrodes vs. four electrodes; steps 8 and 10) in Study 1, deciding thereafter to exclude steps 8 and 9 from Study 2 (presentation of relevant calculations in Results below).

Referring to Study 1, we controlled for the hemispheric lateralization after measuring and analyzing the whole scalp alpha-band activation as projected on all available electrodes and 100% of the EEG data collected for each composition and subject (Step 8). Additionally, looking to validate the lateralized alpha-band activation further, a major existing dipoles calculation was also performed using the BESA model (MEGIS Software GmbH, Munich) (Step 9). This step revealed the most active dipoles (100% of the EEG data) for each musical composition and subject, providing an extra level of evaluation of the hemispheric lateralization of continuous-response activation. At a second level, we focused only on the closest to the four main electrodes, the Smidt and Trainor (2001) study used before.

After statistically validating our dependent variables, our calculations further followed a two-level factor analysis. These factors were presented as ‘between-subjects’ and ‘within-subjects’ codes. For the ‘between-subjects’ code, we compared the continuous biological evoked response of emotion between subjects, nominating it to be the macro-level response. For the ‘within-subject’ code, we attempted to showcase the inherent trajectory of the continuous biological evoked response of emotion for each participant and individual musical composition, respectively. This time, this code was nominated to be the micro-level response.

Based on this two-level factor analysis, we used a Mixed Design ANOVA model (Fig 4) on the evoked and measured alpha brainwaves on each tag point, computing

a. the statistical significance of the dominant hemispheric activation for each tag and composition and for each participant separately (within-subject analysis; repeated measures ANOVA) in order to study the cohesion of emotional biological responses (biological quality of micro-responses); and
b. the statistical significance of the overall dominant hemispheric activation for each composition and for each participant in cross-comparison (between-subject analysis; independent ANOVA) to study the cohesion of emotional biological responses for each musical composition at a combined sample level (biological quality of macro-responses of emotion).

**Fig 4.**
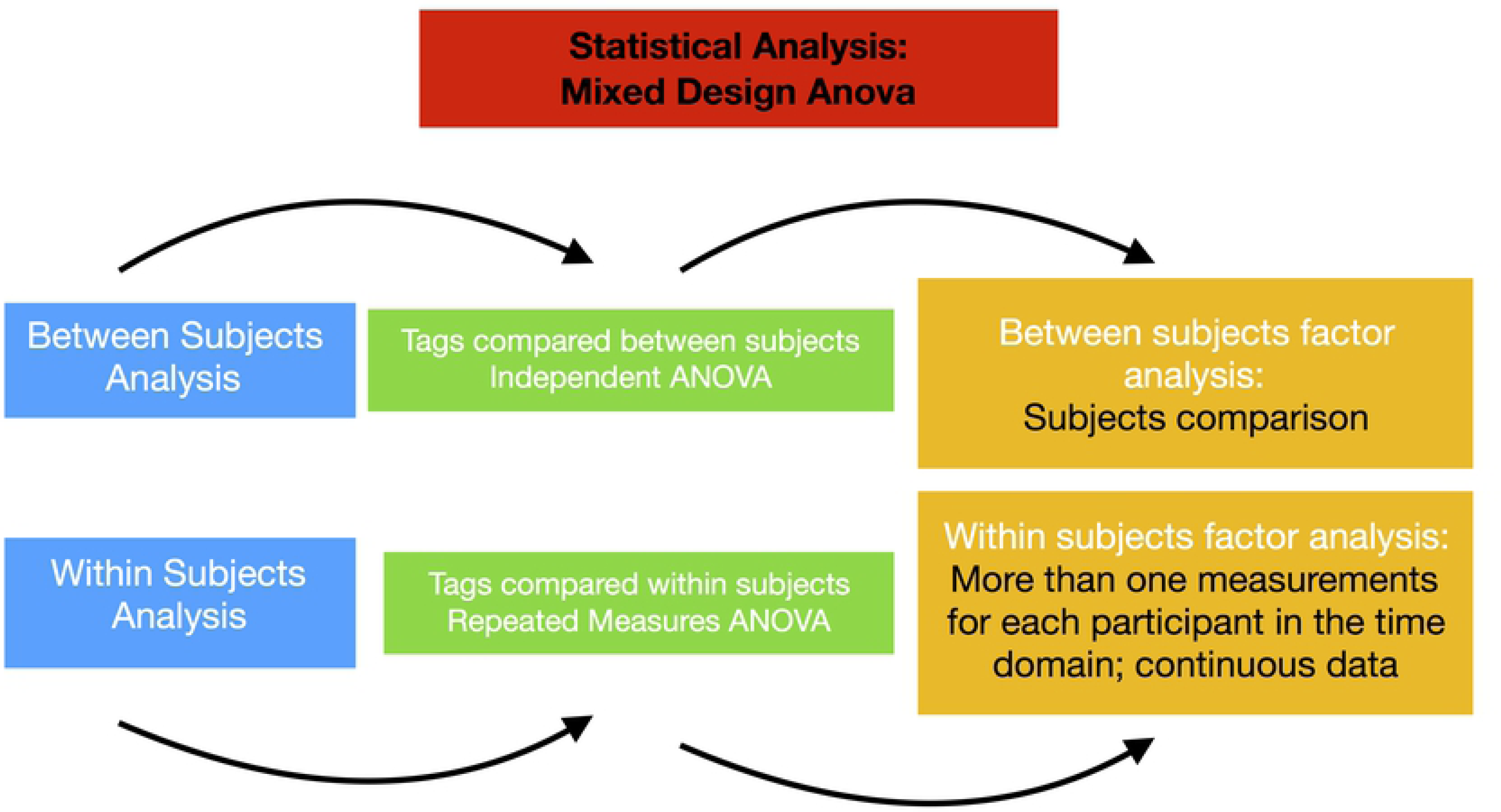
Overview of the statistical analysis

For the aforementioned first case, we followed the approach of the repeated measures ANOVA in order to see how much of the variability shown is a result of the experimental manipulation (tag points in musical stimuli), relative to other random factors (residual) existing in our data collection process (for example, exogenous sound stimulation, prior emotional dynamics etc.) while in the second case, we followed the approach of the independent ANOVA in order to establish if variance differences between participants are isolated or not. In this latter case, the resulting F-test for structuring a sufficient baseline for the overall emotional, biological response for each musical composition can be more potent at a combined sample level.

## Results

In order to categorize our sample in advance on possible differentiations and variances due to brain structure qualities, we ran an audiometric threshold test using the Auditory Brainstem Response (ABR) technique, as well as a Posterior Dominant Rhythm (PDR) analysis. As far as the ABR analysis is concerned, all subjects had normal wave V click-evoked latencies (<6.8 ms in response to the parameters set on structuring the ABR), both before and after the presentation of the protocol stimuli for both Studies. This test verified that no neurological acoustical disorder existed in any of our participants. The PDR data analysis for all participants also showed no significant variations.

### All electrodes hemispheric lateralization

#### Study 1

The immediately following results (Table 2) refer to the 128 electrodes 100% full scalp data, focusing on the alpha band hemispheric lateralization for all three subjects recorded in Study 1. R (right) and L (left) present the corresponding dominating hemispheric alpha band activation on each tag, for each different musical composition for each participant. These calculations were based on the alpha band frequencies spectrum analysis (8Hz; 10Hz; 12Hz) provided through calculated spectoplots for each tag and participant separately (for example, Figs 5 and 6).

**Fig 5.**
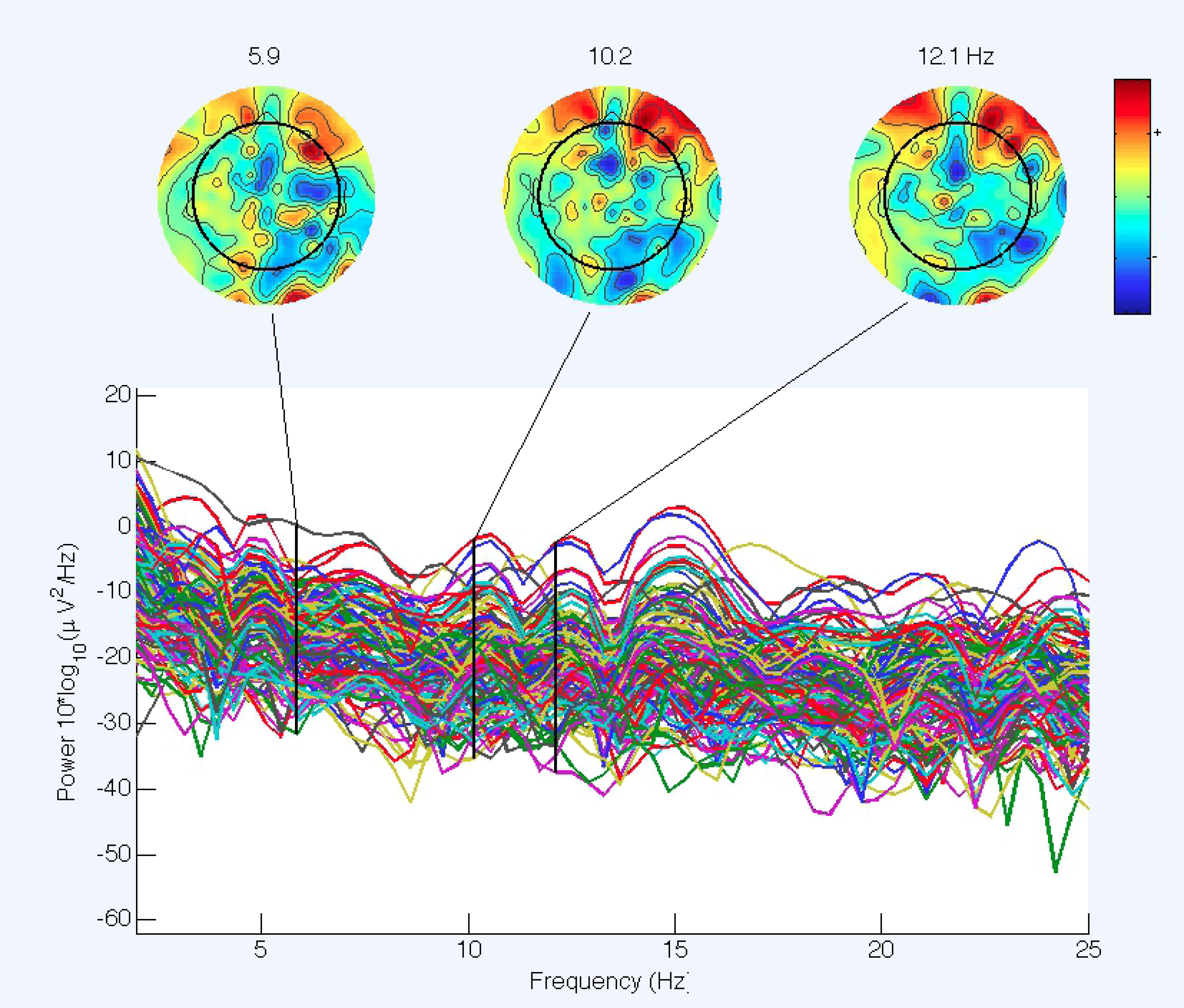
Example of Subject 3, S8 Tag full scalp spectoplot alpha band; 100% of data for S8 tag epoch; 1 sec baseline-2 secs active response.

**Fig 6.**
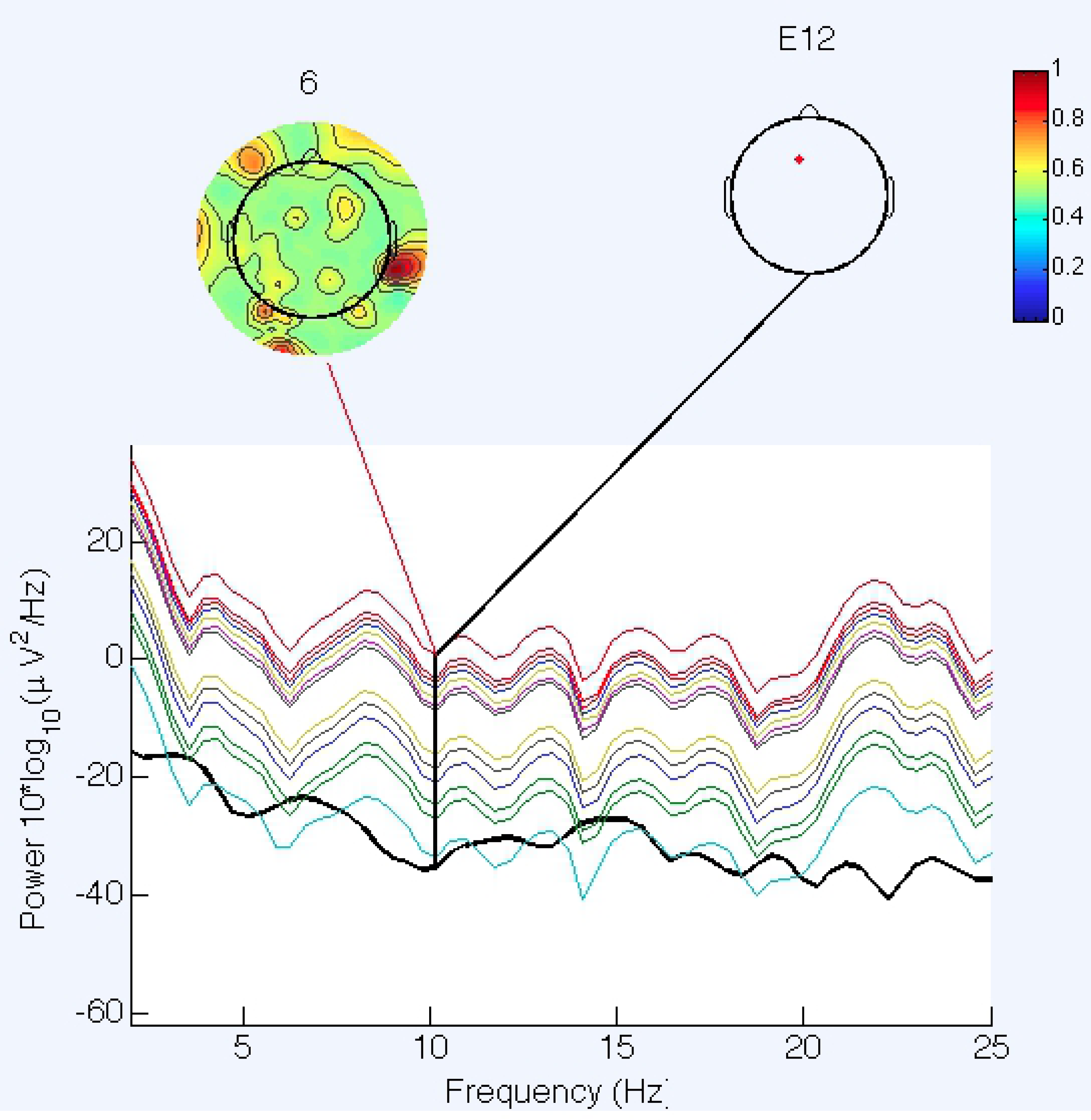
Example for Subject 3, S8 Tag, dominant component lateralization on the electrode with maximum power in the specific epoch.

**Table 2.**
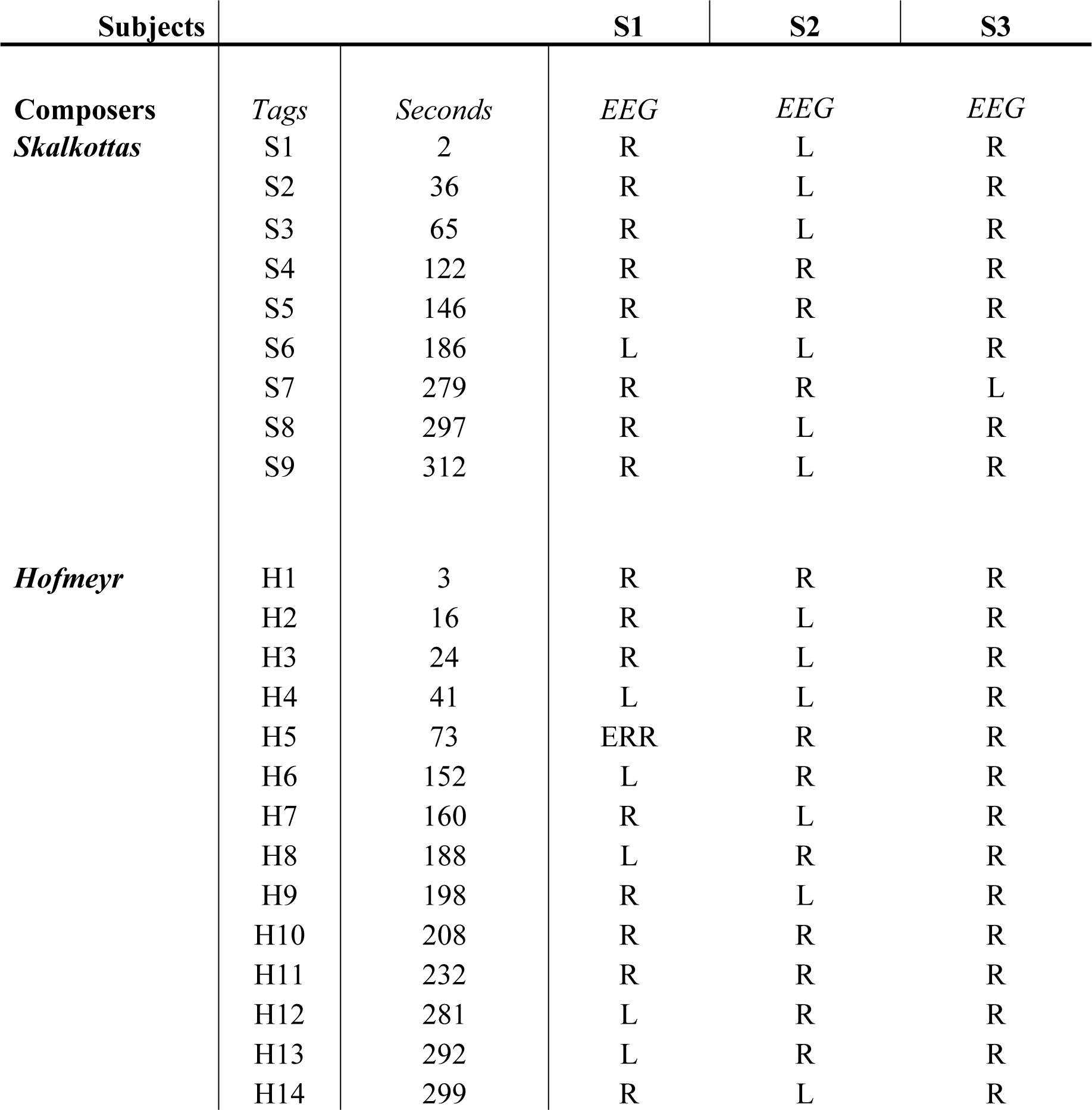
Dominating hemispheric alpha band activation on each tag, 128 electrodes. L = Left; R = Right; ERR in EEG = could not be calculated.

| Subjects |  |  | S1 | S2 | S3 |
| --- | --- | --- | --- | --- | --- |
| Composers | Tags | Seconds | EEG | EEG | EEG |
| *Skalkottas* | S1 | 2 | R | L | R |
|  | S2 | 36 | R | L | R |
|  | S3 | 65 | R | L | R |
|  | S4 | 122 | R | R | R |
|  | S5 | 146 | R | R | R |
|  | S6 | 186 | L | L | R |
|  | S7 | 279 | R | R | L |
|  | S8 | 297 | R | L | R |
|  | S9 | 312 | R | L | R |
| *Hofmeyr* | H1 | 3 | R | R | R |
|  | H2 | 16 | R | L | R |
|  | H3 | 24 | R | L | R |
|  | H4 | 41 | L | L | R |
|  | H5 | 73 | ERR | R | R |
|  | H6 | 152 | L | R | R |
|  | H7 | 160 | R | L | R |
|  | H8 | 188 | L | R | R |
|  | H9 | 198 | R | L | R |
|  | H10 | 208 | R | R | R |
|  | H11 | 232 | R | R | R |
|  | H12 | 281 | L | R | R |
|  | H13 | 292 | L | R | R |
|  | H14 | 299 | R | L | R |

The data in Table 2 were also verified following a major dipoles analysis on 100% of the data for each one of the tags of the two musical compositions. Using the BESA model of analysis software (MEGIS Software GmbH, Munich) as embedded in the EEGLab, we computed the existing major dipoles concentration for each participant and each musical composition individually. Again, we focused on the same band of activation (for example, Fig 7).

**Fig 7.**
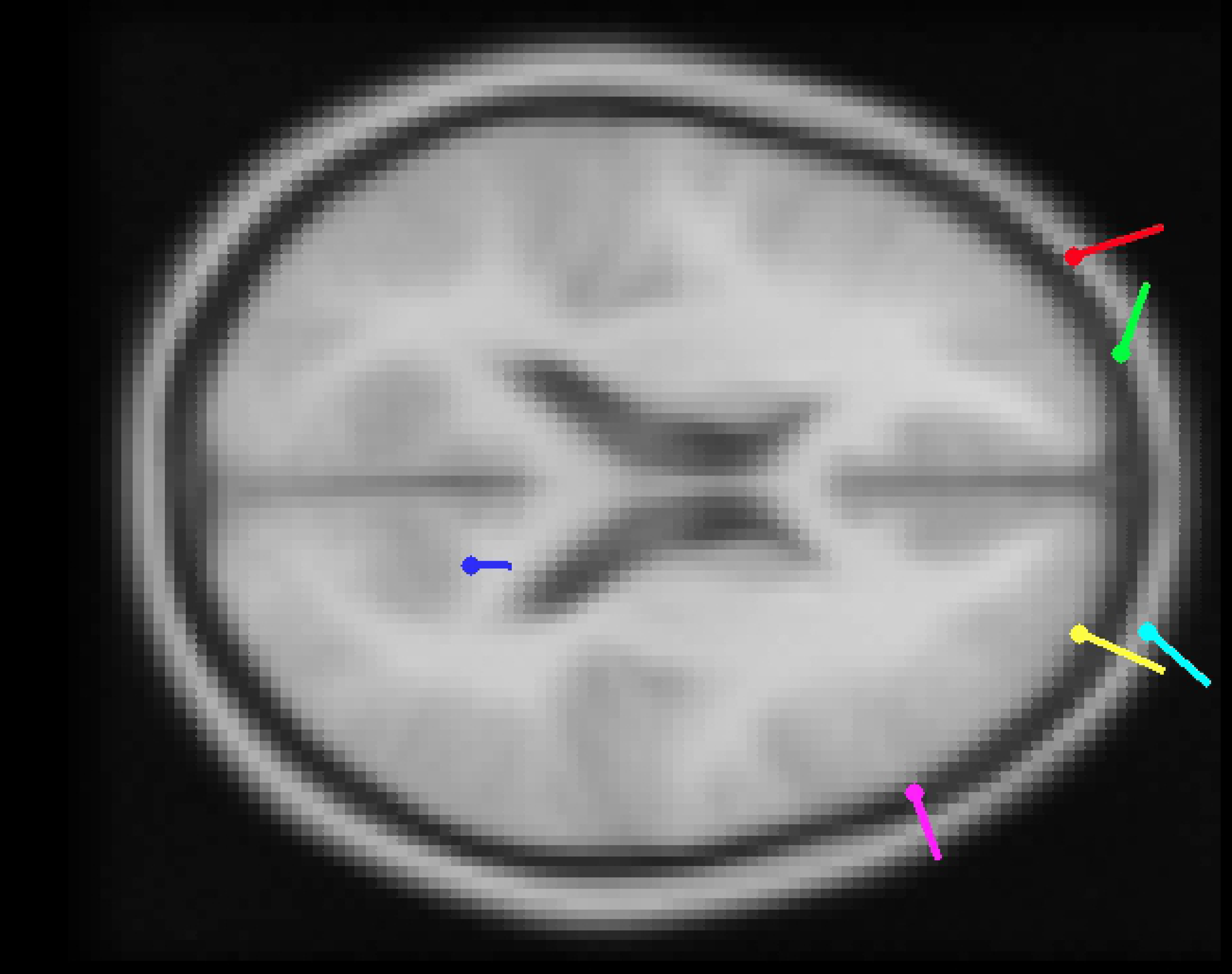
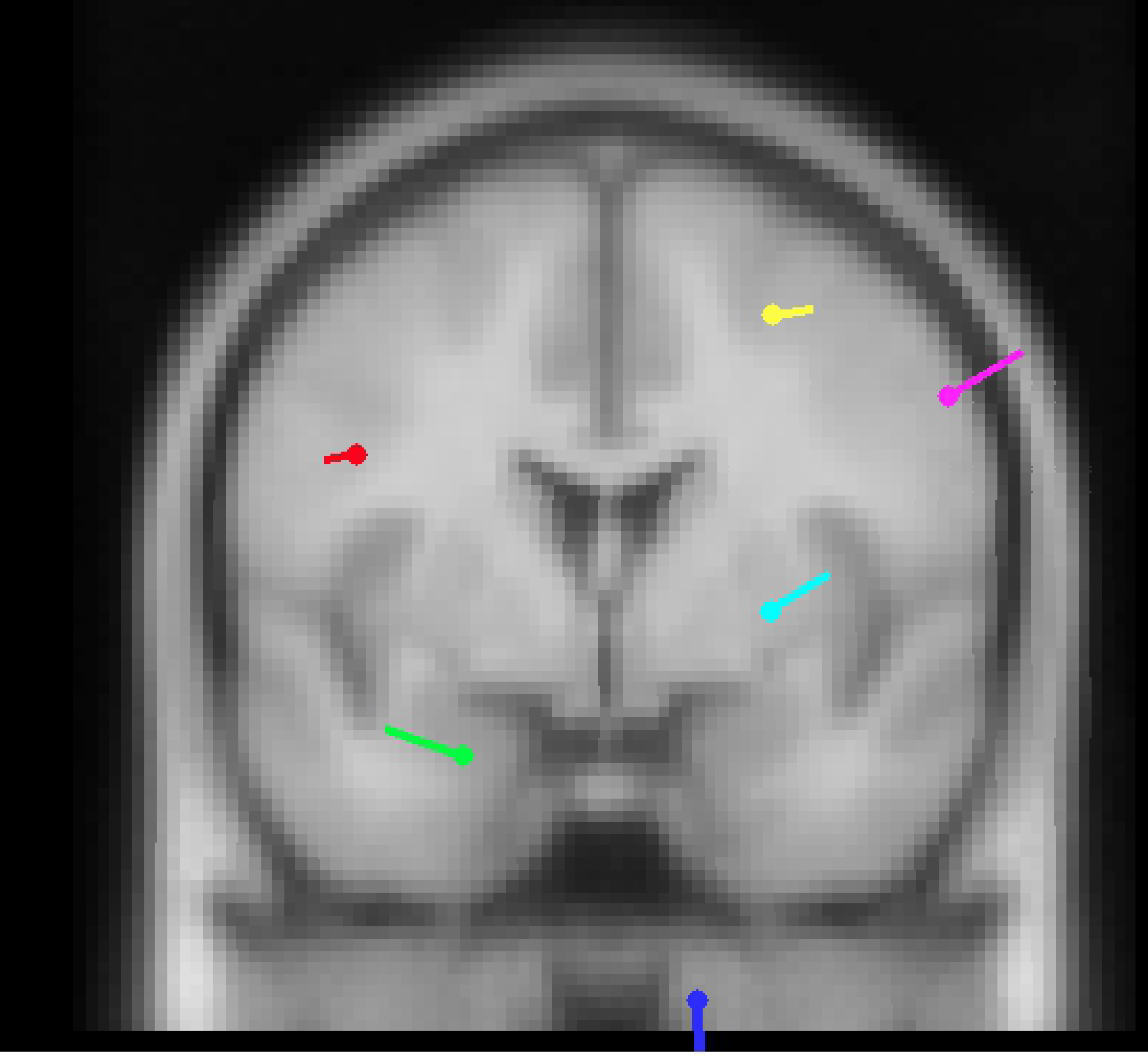
Subject 3, Tag S8 Major Dipoles concentration in 100% of epoched data; Averaged alpha power

### Four electrodes hemispheric lateralization

#### Study 1

We investigated the respective alpha band asymmetric activation between hemispheres for each subject and musical composition separately. We first computed the alpha power for the F3, F4, P3 and P4 electrodes as corresponding to the 128 EGI electrodes system system (electrode numbers 25, 124, 53 and 87) [71] continuing with a computation of the activation asymmetry ratios between the pairs of these four electrodes according to the hemispheric index formula (var(x) – var (y)) / var (x) + var (y)) that has been reliably used in previous similar investigations (for example [72, 73]). In this case (x) and (y) equals to channel variance index of the filtered output on the left side and the right side, respectively.

On the tag point number S8 for Subject 3, for example, calculations on the two pairs of electrodes (25:124 and 53:87) rendered the following index numbers: for the 25:124 pair of electrodes, anteriorly lateralized asymmetry computes at 0.4703 while for the 53:87 pair, posteriorly lateralized asymmetry computes at −0.5882. These numbers mean that we got an overall positive left side anteriorly localized domination of the alpha band for this set of electrodes. Equally, for all the available pairs of electrodes and data, we computed asymmetry reaching the result shown in Table 3 below. Statistically comparing the results of Table 3 to Table 2 above (paired t-tests), we calculated the mean difference between the two sets of measurements (whole set of electrodes vs. four electrodes). We found no statistically significant differences between the two pairs of measurements (The p-values for each pair was: [S1 Skalkotas p=1, S1 Hofmeyr p=0.605; S2 Skalkotas p=0.5, S2 Hofmeyr p=-0.043; S3 Skalkotas p=0.661, S3 Hofmeyr p=1).

**Table 3.**
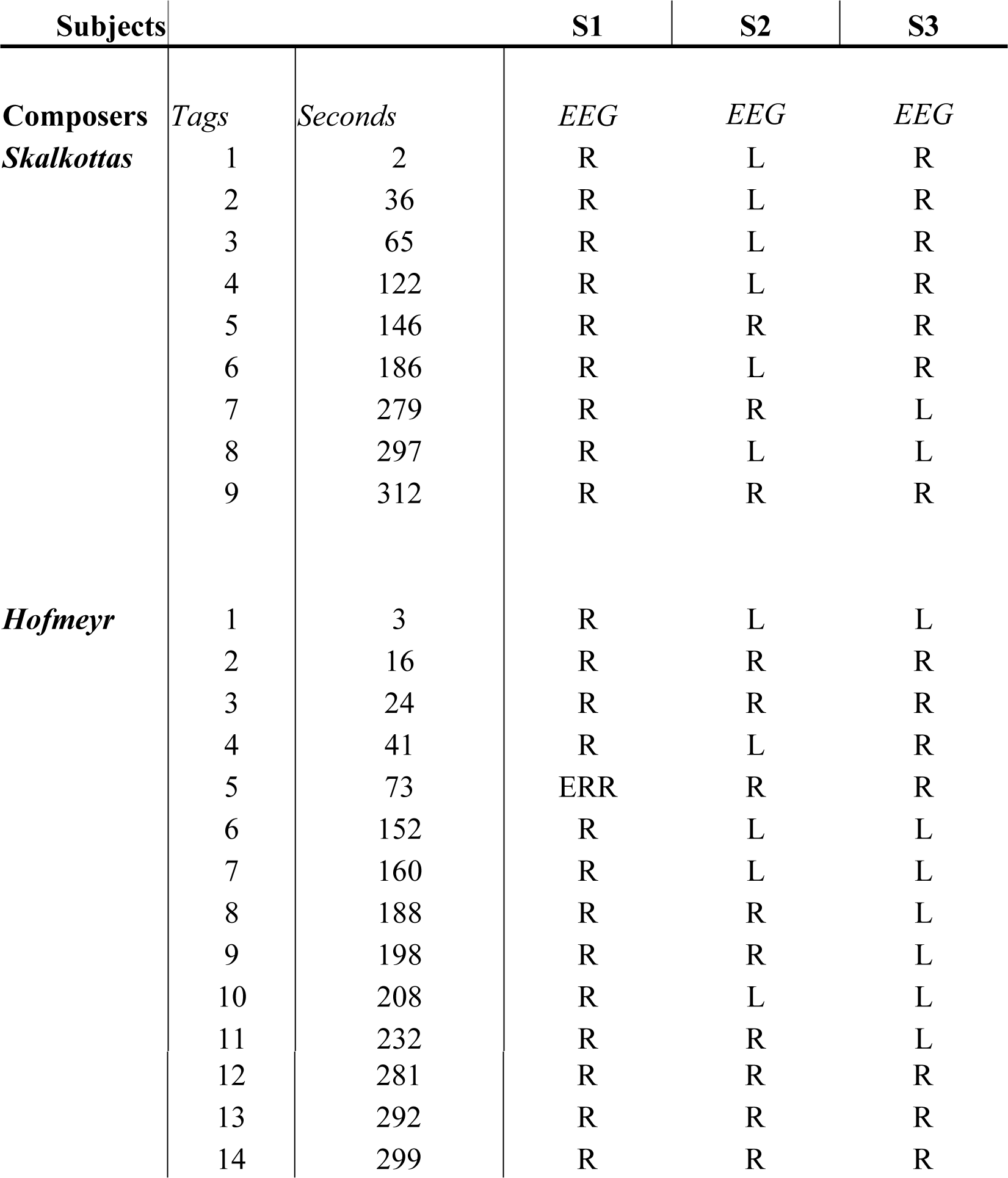
Dominating hemispheric alpha band activation on each tag, 4 electrodes, in Study 1. L = Left; R = Right, ERR in EEG = could not be calculated

#### Study 2

We investigated the respective alpha band asymmetric activation between hemispheres for each subject and musical composition separately. We first computed the alpha power for the F3, F4, P7 and P8 electrodes as corresponding to the 14 Emotiv+ EPOC unit electrodes system, continuing with a computation of the activation asymmetry ratios between the pairs of these four electrodes according to the hemispheric index formula (var(x) – var (y)) / var (x) + var (y)) as employed in Study 1 before. For all the available pairs of electrodes and data, we computed the following asymmetry results (Table 4).

**Table 4.**
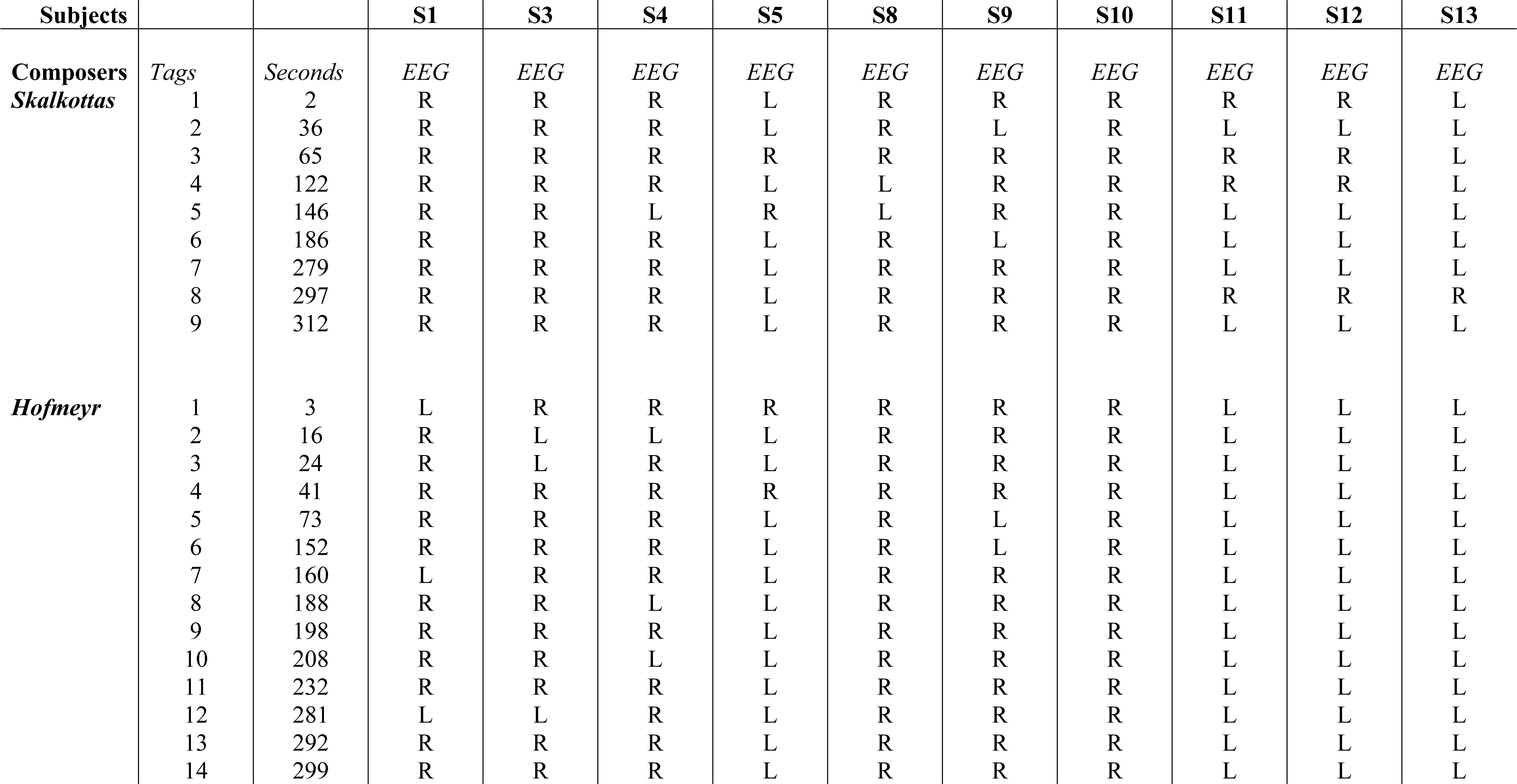
Dominating hemispheric alpha band activation on each tag, 4 electrodes, Study 2. L = Left; R = Right.

### Factor analysis

With these two studies, we aimed to investigate the micro- and macro-emotional responses following an intra-subjective as well as inter-subjective data analysis. Therefore, two specific hypotheses supported the statistical analyses of the neurophysiological data. The intra-subjective statistical analysis was based on the null hypothesis (H0_1_), suggesting that all tags in an emotionally organized longitudinal sound stimulus evoke a solid emotional soundscape. With this in mind, we followed for the two main conditions (the two musical compositions) a repeated-measures one-way ANOVA (RM-ANOVA) on the tags for each musical composition and each subject. On the inter-subjective statistical analysis side, the null hypothesis (H0_2_) we followed, suggested that all subjects present similar or close to similar a biological reaction (hemispheric lateralization) in relation to the tags inserted in the musical composition. In order to test this hypothesis, we followed for the two main conditions (the two musical compositions) an independent ANOVA computation on the tags for each musical composition, yet including the subjects as different variables in the same equation this time, knowing from our previous calculations that indeed, each of these variables perform and stand as uniform (i.e., as continuous and not as fluctuating evoked responses intra-subjectively).

#### Study 1

For the H0_1_, following a significance level of α=0.05 in our calculations and a Bonferroni correction process, the Mauchly’s test indicated first that the assumption of sphericity was not violated in both stimuli measurements. Then, the RM-ANOVA results showed that there was no significant effect on the whole compositions’ evoked emotional responses through any unique representations of the tags’ emotional power as presented to the biological reactions [S Stimulus: *F*(8) = 0.684, *p* = .07] [H Stimulus: F(13) = 0.451, *p* = 0.933]. These findings suggest that each subject reacted at an average level uniformly in terms of hemispheric activation for each different composition. In other words, the emotional carriage, as shown in terms of brain activation, was almost uniform, suggesting that the subjects indeed perceived each musical composition as having an average, holistic emotional load.

For the H0_2_, contrary to the previous results of the intra-subjective approach, the inter-subjective calculations showed that there is a significant effect of the emotional power carried from the different tag points on the biological reactions (hemispheric lateralisation) [S Stimulus: *F*(1) = 84.64, *p* = .012] [H Stimulus: F(1) = 117.23, *p* = 0.008]. These findings suggest that the subjects indeed reacted differently in terms of hemispheric lateralization for each of the different composition’s tag lines. In other words, the emotional progression if shown in biological terms for each musical composition, follows a completely different synthesis for each individual.

#### Study 2

For the H0_1_, following a significance level of α=0.05 again in our calculations and a Bonferroni correction process, the Mauchly’s test indicated that this time, the assumption of sphericity was violated in both stimuli measurements. Therefore, we applied the RM-ANOVA calculations with a Greenhouse-Geisser correction. Through this correction, we determined that the mean brain lateralization did not differ statistically significantly between either the 14 noted tags for the H Stimulus [F(3.944, 35.495)=0.866, p=0.493] or the nine noted tags for the S Stimulus [F(3.339, 30.055)=2.279, p=0.094]. Post-hoc tests for both stimuli, using the Bonferroni correction, revealed that the continuous stimuli of the Hofmeyr and the Skalkotas musical pieces elicited no actual differentiation in terms of brain activity lateralization (p=1). In other words, the emotional carriage, as shown in terms of brain activation for each tag was almost uniform, suggesting that the subjects indeed perceived each musical composition as having an average, holistic emotional load. This result is on par with the results of Study 1.

For the H0_2_, contrary to the HO_2_ of the Study 1, the inter-subjective calculations showed that there is no significant effect of the emotional power carried from the different tag points on the biological reactions (hemispheric literalization) [S Stimulus: *F*(8, 81) = 1.251, *p* = 0.281] [H Stimulus: F(13, 126) = 0.263, *p* = 0.995]. These findings suggest that the subjects reacted similarly in terms of hemispheric lateralization for each of the different composition’s tag lines. In other words, the emotional progression shown in biological terms for each musical composition follows a similar progression line for each individual.

## Conclusion

The above results present the first evidence of our study for the emotionally induced hemispheric lateralized activity as evoked through a continuous listening process of two musical compositions diverse in cultural and musical technical elements. With our protocol, we proposed an inter-subjective as well as an intra-subjective pathway of data analysis, and we respectively studied the assigned musical compositions at a macro- and a micro-emotional level of response considering a non-personalised response framework. After recruiting and investigating two unrelated in terms of musical expertise, cultural and educational background cohorts (i.e. EEG Study 1 and EEG Study 2), we found that there is quite a similar approach for both of them on how they finally shape their emotional reaction at the macro-response level (Study 1 = Study 2). Therefore, we conclude at this initial stage that a musical composition – when seen as a continuous and holistic emotional response – may indeed potentially induce an aligned biological response of emotion, no matter what the differences are in terms of the equipment used to measure these responses or the demographics of the recruited sample.

At the micro-response level, however, our study provided an entirely different picture. In this case, our two studies presented a quite detached profile of statistical differentiation (Study 1 ≠ Study 2) suggesting that any specific structural parts of music evoking the holistic emotional reaction through a continuum, may not biologically be perceived in the same way by individuals encompassing different demographics. It seems that the quality of the biological steps a person follows in order to conclude to a particular emotion on a specific musical composition could be variable, depending on either to the cultural or the educational elements portraying their personal background. This fact proves, on the one hand, our emotions’ complexity as human beings, as well as on the other the intricacy of devising and running relevant to music protocols in the particular field of behavioral neuroscience. It seems that structural elements of music are indeed able to fore-bring a complex footprint of emotional reaction, different for individuals with detached backgrounds and contexts, even if undergoing the same listening experience, whatsoever.

If a fully-understood quality of an emotional response is the ultimate goal, then, the above datum-point begs for a multi-layered investigation approach. This multi-layered approach can be achieved, in our point of view, by a grounded emotional variability framework that could serve as a functional basis of organization and growth in context (i.e., investigation of similarities and differences in context over time). For this reason, we intend to further introduce in our study the micro developmental variation dynamics theorem [74]. According to the theorem, when functional characteristics of human beings studied in reference to emotional development, three units of analysis should be there to conclude the study successfully. These three units are “the intra-individual variability [relative rapid and reversible changes or differentiations], [the] intra-individual changes [relative stable changes or differentiations], and [the] inter-individual differences (highly stable changes over a long-time period or among individuals]” [74]. In our study, it seems that we have already incorporated the ‘intra-individual variability’ and the ‘inter-individual differences’ units when measuring the intra-subjective and inter-subjective factors. What remains is to integrate the ‘intra-individual changes’ unit further as required, measuring emotional reactions to a particular musical piece more than once over different timeframes (for more details on this possible integration, please read [75]).

Concluding, the next steps will indeed follow a more extended framework of analysis through the micro developmental variation dynamics theorem, as well as further steps to recruit a more extended pool of participants and stimuli to investigate and understand at a biological level. Finally, we will conclude our intended biopsychological analysis and comparison, when both the neurophysiological as well as the behavioral data will be statistically and qualitatively combined. Bio-psychologically measured emotional reactions to whole musical compositions is definitely an under-researched subject in the field of behavioral neuroscience, and therefore worthing of the most detailed and robust possible basic research; especially when someone considers their full potential in real-life applications which certainly extends to more than the clinical rehabilitation settings (e.g., Alzheimer’s disease) or the socioemotional regulation applications through music (e.g. depression).

## Acknowledgments

We would like to acknowledge the help and advice given by Theo Herbst, Barry Ross, Miles Warrington, and Vasiliki Pliogou in actualizing and completing this study successfully.

## Footnotes

i Music can be defined as ‘sounds that are subject to some form of human organization (either in production or reception or both) and communication that is not the same as spoken or written language’, with ‘internal affective processing’ as one of the contributing factors to the universality of music (please see the views of Agawu [76] on the myth that music is a universal language. He makes it very clear that although music shares some properties of language, it cannot be considered as a language in a linguistic sense.

ii The field of Musicology has been traditionally subdivided into History Musicology and Systematic Musicology (1885) [78]. In this division Historic Musicology deals with research related to historic issues, and Systematic Musicology with music-theoretical, structural and style-related formal issues. Since Adler’s two-tiered division, the field of Musicology has expanded into numerous subdisciplines, of which Neuromusicology is but one of a multitude of fields that are researched [79]. Despite the addition and rearrangement of subdisciplines in the overarching umbrella term, Systematic Musicology still refers to the study of music theoretic principles. The music-theoretical analyses methods have been standardised. As such there is no need for music-theoretical analyses of the two compositions by a panel of experts. The music-theoretical tools that are used in the analyses are universal, at least for Western Classical Music, as represented in the two different musical compositions.

